# Structural and histone binding studies of the chromo barrel domain of TIP60

**DOI:** 10.1101/257485

**Authors:** Yuzhe Zhang, Ming Lei, Xiao Liang, Peter Loppnau, Yanjun Li, Jinrong Min, Yanli Liu

**Author notes:** These authors contributed equally to this work (Yuzhe Zhang, Ming Lei).

## Abstract

TIP60 consists of an N-terminal chromo barrel domain (TIP60-CB) and a C-terminal acetyltransferase domain and acetylates histone and non-histone proteins within diverse cellular processes. Whereas the TIP60-CB is thought to recognize histone tails, molecular details of this interaction remain unclear. Here we attempted a quantitative analysis of the interaction between the TIP60-CB and histone peptides, but did not observe any binding through either fluorescence polarization or isothermal titration calorimetry. We solved a crystal structure of the TIP60-CB alone. Analysis of the crystal structure demonstrates a putative peptide binding site that may be occluded by the basic side chain of a residue in a unique β hairpin between the two N-terminal strands of the β barrel.

## Introduction

TIP60 (tat-interactive protein 60), also known as KAT5 or HTATIP, is a MYST (MOZ, YBF2, SAS2 and TIP60) family acetyltransferase that comprises an N-terminal chromo barrel domain (TIP60-CB) and a C-terminal histone acetyltransferase domain[1–3]. TIP60 catalyzes acetylation of lysine side chains in various histone[4,5] and non-histone[6,7] proteins, including itself[8], and plays critical roles in multiple cellular processes, such as chromatin remodeling[5], transcription[5], DNA double-strand break (DSB) repair[3,9], apoptosis[9], embryonic stem cell identity[10] and embryonic development[11]. Abnormal expression of TIP60 is associated with tumorigenesis of several carcinomas, such as lymphomas, head-and-neck, breast[12] and prostate cancers[13]. Homozygous loss of *Tip60* leads to embryonic lethality in mice[11].

Several studies have analyzed the interaction between modified histones and TIP60-CB. It was found that TIP60-CB recognizes trimethylated lysine at site 9 of histone H3 (H3K9me3), which triggers TIP60 to acetylate and activate ATM, thereby promoting the DSB repair pathway[3]. Phosphorylation of tyrosine 44 of TIP60 reportedly strengthens the interaction between TIP60-CB and H3K9me3 and enhances the activity of TIP60, again facilitating the DSB repair pathway[2]. According to another study, TIP60-CB interacts with monomethylated lysine at site 4 of histone H3 (H3K4me1), which stabilizes TIP60 recruitment to a subset of estrogen receptor alpha (ERα) target genes, and then facilitates regulation of the associated gene transcription[1].

Based on the published reports of histone-TIP60 binding, and the demonstrated binding between histones and TIP60 homologs, we aimed at characterizing the interaction between TIP60-CB and histone peptides by quantitative binding assays and crystallographic analysis. In this study, we tried to analyze the interactions between TIP60-CB and histone peptides by quantitative fluorescence polarization (FP) and isothermal titration calorimetry (ITC), however we could not detect any obvious binding. We attempted to co-crystallize TIP60-CB with its reported partners and solved the peptide unbound structures only. Analysis of the TIP60-CB apo-structure and comparison with other published chromo barrel structures reveal possible reasons for our failure to detect histone binding in solution or to determine a co-crystal structure of the TIP60-CB in complex with a histone peptide.

## Results and discussion

### Interaction between histone peptides and TIP60-CB not detected by ITC or FP assays

According to previous reports, TIP60-CB binds to H3K9me3 peptide[3]; binding is enhanced by phosphorylation at Tyr44 of TIP60[2]. Our own isothermal calorimetry (ITC) assays failed to detect such binding to the H3K9me3 peptide with or without tyrosine kinase treatment of TIP60 (Fig. 1A and 1B). TIP60-CB reportedly also binds to H3K4me1[1], which we again failed to detect (Fig. 1C). Furthermore, we screened a library of peptides containing the major methylation modifications on histones H3 and H4, including H3K4/9/27/36/79me0-me3, H4K20me0-me3 and H3R2 mono-, symmetric or asymmetric di-methylarginine, aginst TIP60-CB by fluorescence polarization (FP) assay (data not shown), but did not detect any significant interactions. Although our data are in contrast with the studies mentioned above, they are consistent with a previous protein-domain microarray assay results that failed to detect any TIP60-CB binding to methylated histone H3 or H4 peptides[14]. There are reports of weak (millimolar) peptide binding to proteins such as RBBP1[15], AtMRG1/2[16] and ScEaf3[17], which might fall below our own assays’ limits of detection. Moreover, the TIP60-CB may depend on an as yet undiscovered modification pattern.

**Figure 1.**
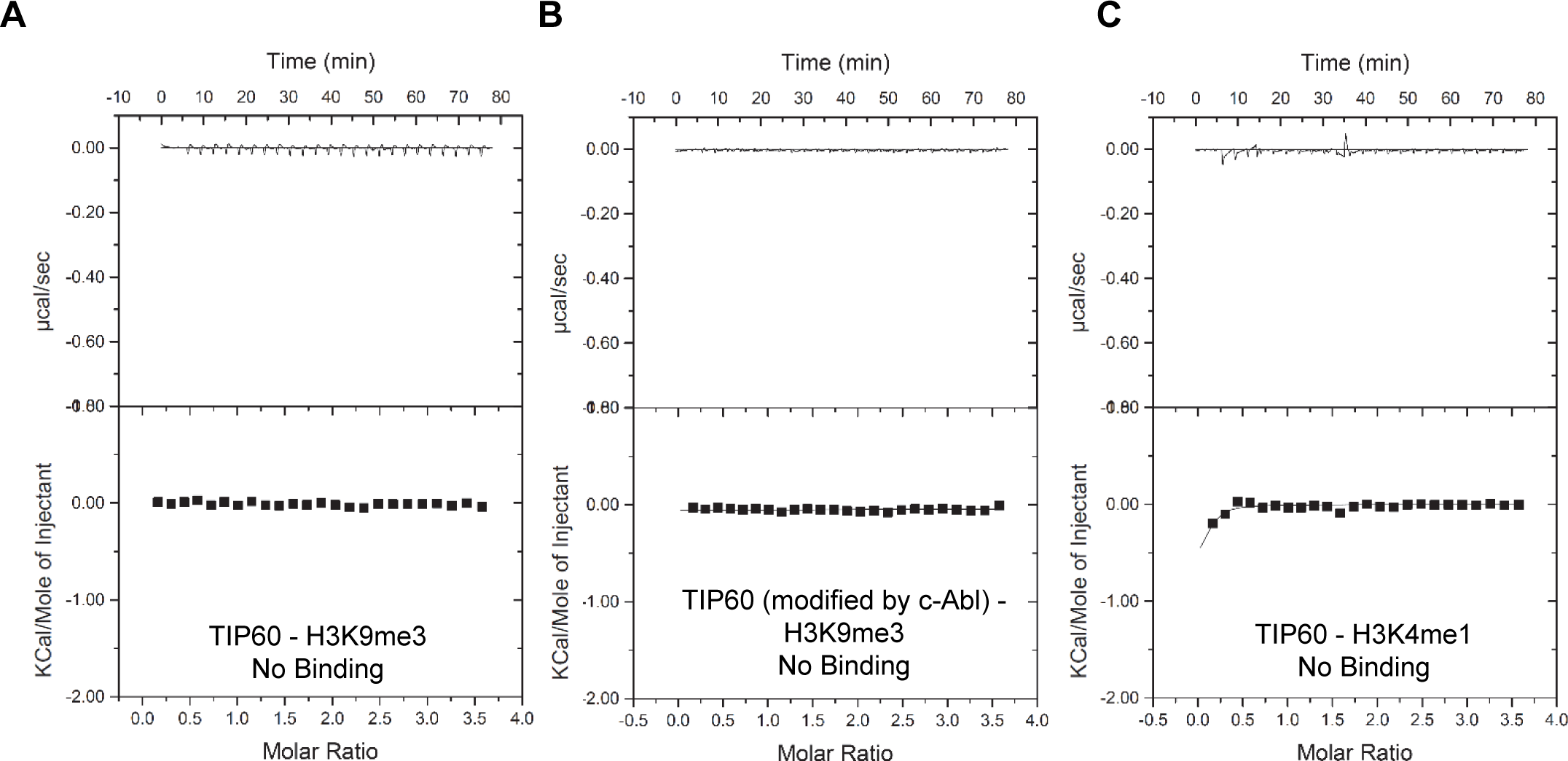
ITC binding curves for titration of different histone peptides to TIP60-CB. **(A and B)** ITC binding curves of histone H3K9me3 peptide to TIP60-CB with (B) or without pre-treating by tyrosine kinase c-Abl (A). **(C)** ITC binding curve of histone H3K4me1 peptide to TIP60-CB.

### TIP60-CB residues Tyr47 and Phe50 may form an aromatic cage

Even though Momen *et al.* have released a solution structure of the TIP60 chromo barrel domain under PDB code 2EKO, we have solved the crystal structure of the chromo barrel domain of TIP60 (residues 1-80, Table 1 and Fig. 2) as a result of our unsuccessful attempts at preparing a crystal of the domain in complex with a histone peptide. The asymmetric unit of our crystallographic model accommodates seven TIP60 chromo barrel domain molecules and each molecule is highly similar to the solution structure. As expected, coordinates at the termini of the solution structure of TIP60-CB (Momen *et al.*, PDB entry 2EKO) vary significantly between the models of the NMR ensemble. However, coordinates of the loop between the two most C-terminal barrel strands are also distributed broadly across the ensemble, which may have functional implications (Fig. 2A). This loop includes residue Phe50, which corresponds to residue Trp382 inside the second MBT repeat of the known histone binder L3MBTL1 (Fig. 2A). Therein, together with Phe379 and Tyr386, Trp382 forms the aromatic cage that recognizes dimethylated Lys20 of histone H4[18]. Aromatic cages are formed by between 2 and 4 aromatic residues and are a characteristic feature of methylation effector proteins that recognize methylated lysine[19,20], methylated arginine[21,22] or even N^6^-methyladenosine[23]. Phe379 of L3MBTL1 can be aligned[24] with Tyr47 of TIP60-CB, but the distance between the Cα atoms of Phe379 and Trp382 of L3MBTL1 lies outside and below the 6.1-9.0 Â distribution of inter-Cα distances between Tyr47 and Phe50 of TIP60 in the NMR model ensemble (Fig 2B). Significant “loosening” of the putative cage should affect the capability of TIP60-CB to recognize histone peptides, but the wide distribution of distances raises the possibility that even smaller or larger distances are conformationally accessible to TIP60. Like the NMR solution structure (Momen *et al.*, 2007), the protomers of the TIP60-CB crystal structure largely resemble the prototypical DmMOF chromo barrel[25] (Fig. 2C). Electron density resolved residues Tyr47 and Phe50 with varying clarity across the seven protomers (Fig. 2D and 2E) and suggested that the loop could adopt a conformation similar to the corresponding loop in the second MBT repeat of L3MBTL1, which carries cage residues Phe379 and Trp382.

**Table 1.**
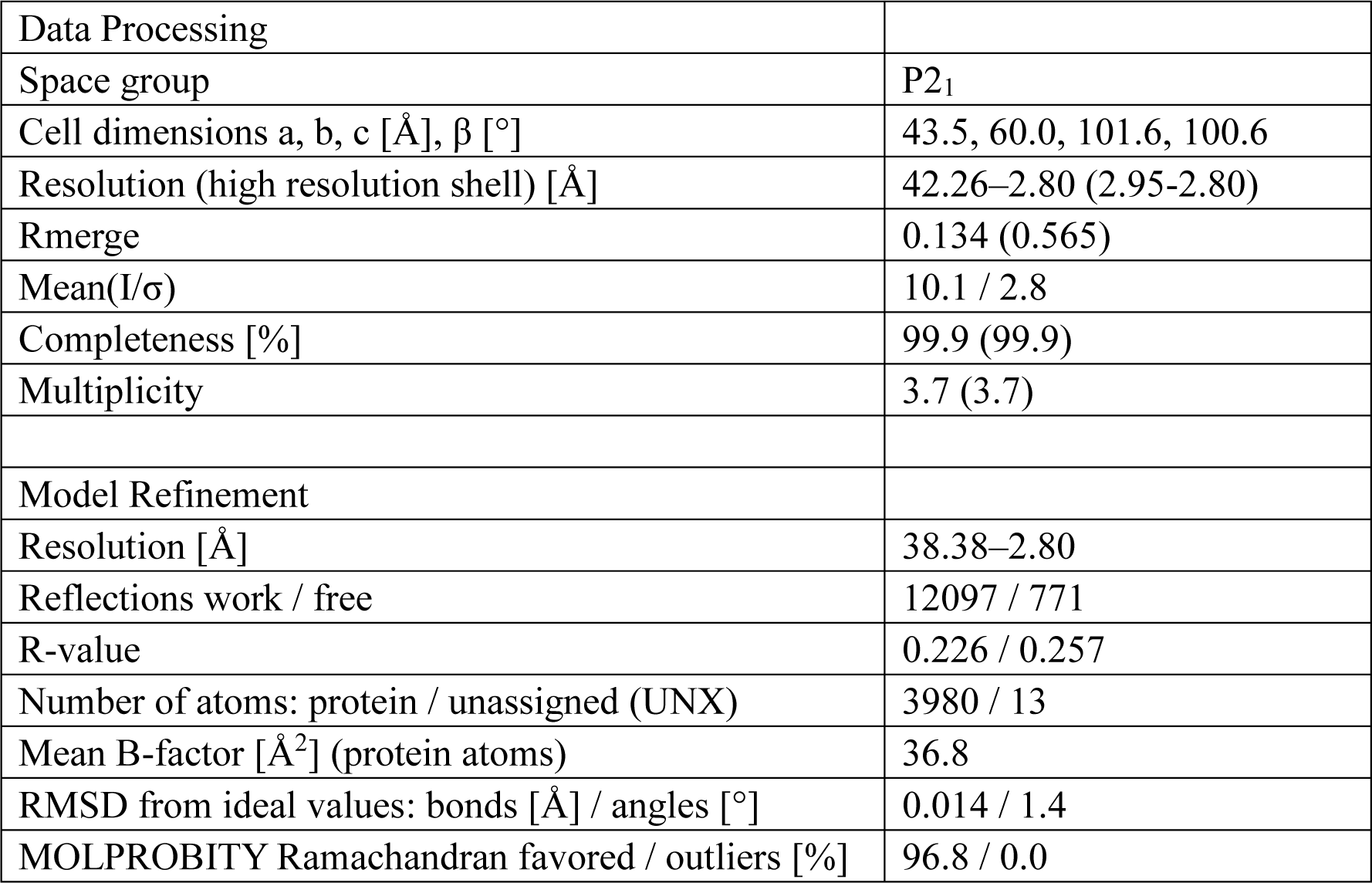
Data collection and refinement statistics

**Figure 2.**
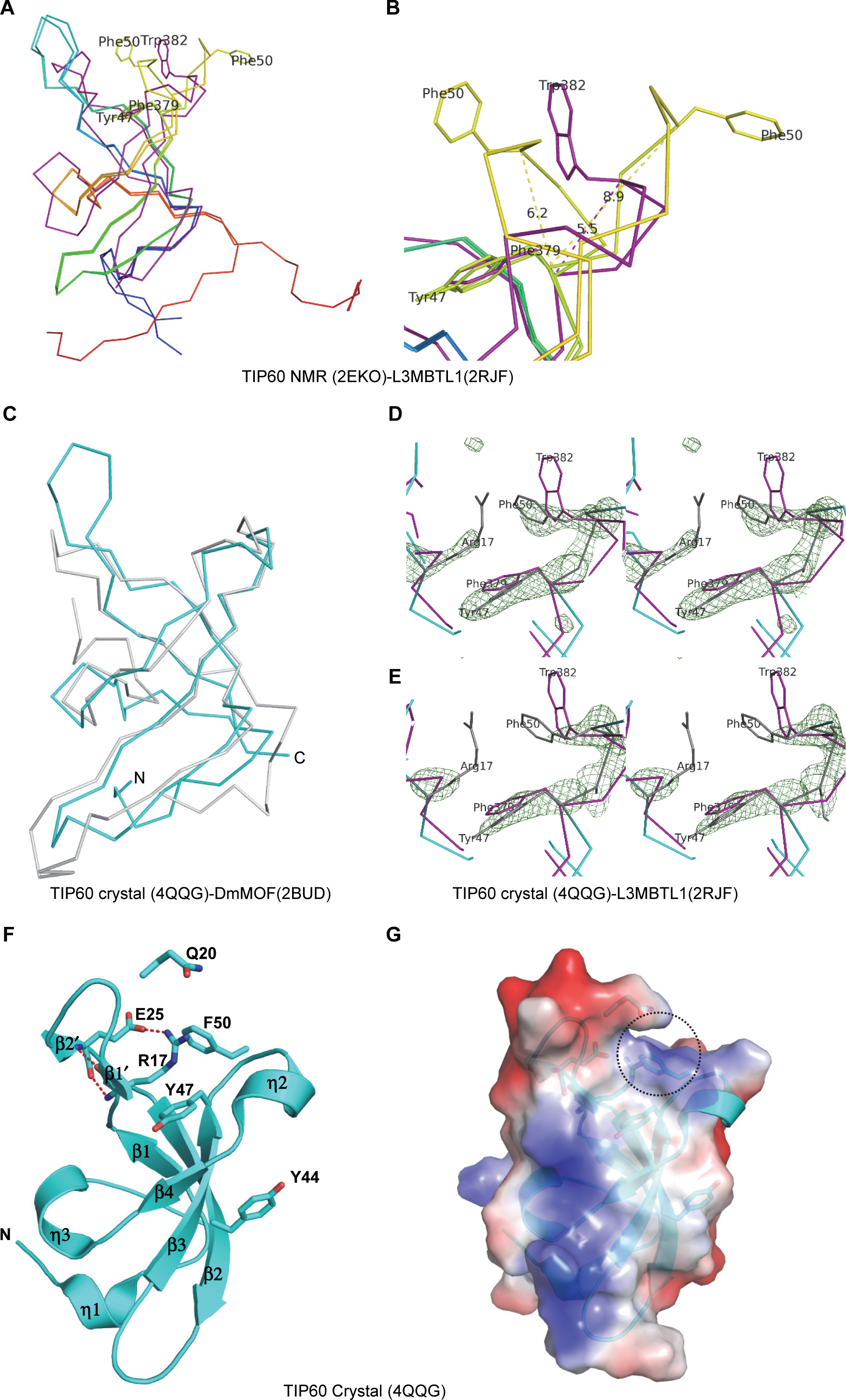
Structure of TIP60-CB. **(A)** Uncertainty in the atomic positions of putative aromatic cage residue Phe50 of TIP60 (numbered Phe53 in PDB entry 2EKO, Momen *et al.).* Two models of the NMR ensemble are rainbow-colored from blue N-terminus to red C-terminus. L3MBTL1 MBT2 (from PDB entry 2RJF) was aligned using the PYMOL implementation of CEALIGN [24] and is shown in purple, including MBT2 aromatic cage residues Phe379 and Trp382. **(B)** Close-up view of two models from the NMR ensemble of TIP60 at the site of the putative aromatic cage. Dashed lines indicate inter-Cα distances. **(C)** Superimposition of TIP60-CB crystal structure (PDB: 4QQG, this work, cyan) and DmMOF (PDB code: 2BUD, gray). **(D)** PHENIX mFo-DFc omit map for TIP 60 residues Arg17 and Tyr47 through Phe50 contoured in PyMOL at level 3 as a green mesh. A Calpha trace of TIP60 is colored cyan, omitted parts of the model are colored gray. The side chain of TIP60 Arg17 is not fully resolved. Also shown, in purple, are Calpha trace as well as Phe379 and Trp382 sidechains of L3MBTL1 (aligned coordinates from PDB entry 2RJF). **(E)** A view of mFo-DFc omit density at level 3 for a different protomer, where the side chain of TIP60 Phe50 is not resolved. Colors and labels correspond to the figure 2B. **(F)** Overall structure of TIP60-CB. The secondary structure elements are shown as cartoons and colored cyan, with the potential cage associated residues shown as sticks. Hydrogen bonds and salt bridges are shown as red dashes. **(G)** Electrostatic potential surface representation of TIP60-CB (isocontour values of ± 76.9 kT/e). Negative and positive potentials are depicted in red and blue, respectively. The potential binding cage resign for the methylated lysine is shown by black circle. Structure figures were generated by using PyMOL (http://pymol.sourceforge.net). Electrostatic potential surface representations were calculated with PyMOL’s built-in *protein contact potential* function[55].

### A possible structural impediment to the binding of the TIP60-CB to lysine-methylated histone peptides

The TIP60-CB, with 4 β-strands followed by a short 310 a-helix, is different from the typical chromo barrel domain comprising a five strands β-barrel and a long α-helix or a short 310 a-helix following these β-strands, such as ScEaf3[17], MRG15[26], MSL3[27], and DmMOF[25] (Fig. 2F and 3A). In addition, TIP60 has a unique β hairpin between the first and second β-strands (Fig. 3B and 3C). As mentioned above, the aromatic residues Tyr47 and Phe50 of TIP60-CB may form an aromatic cage, however, a positively charged side chain of Arg17, locating in the unique β hairpin, occupies the cage (Fig. 2F). The Arg17 side chain is not well resolved in the crystal structure (Fig. 2D and 2E), but residual difference density indicates that the side chain may compete with a putative histone ligand for space in the aromatic cage (Fig. 3D). The residue Arg17 can be readily aligned with residue His21 in the MRG15 chromo barrel domain (Fig. 3B), which has been observed as the obstruction of the aromatic cage by its side chain previously[26]. In addition, the β hairpin, including a long side chain of Gln20, blocks the entrance of the aromatic cage (Fig. 2G and 3D). As binding of methylated histone peptides has nevertheless been demonstrated for the chromo barrel domains of MRG15[26] and its yeast homology ScEaf3[17], such binding can also not be ruled out for TIP60-CB, but would likely entail significant movement of the hairpin between the two N-terminal strands of the chromo barrel.

**Figure 3.**
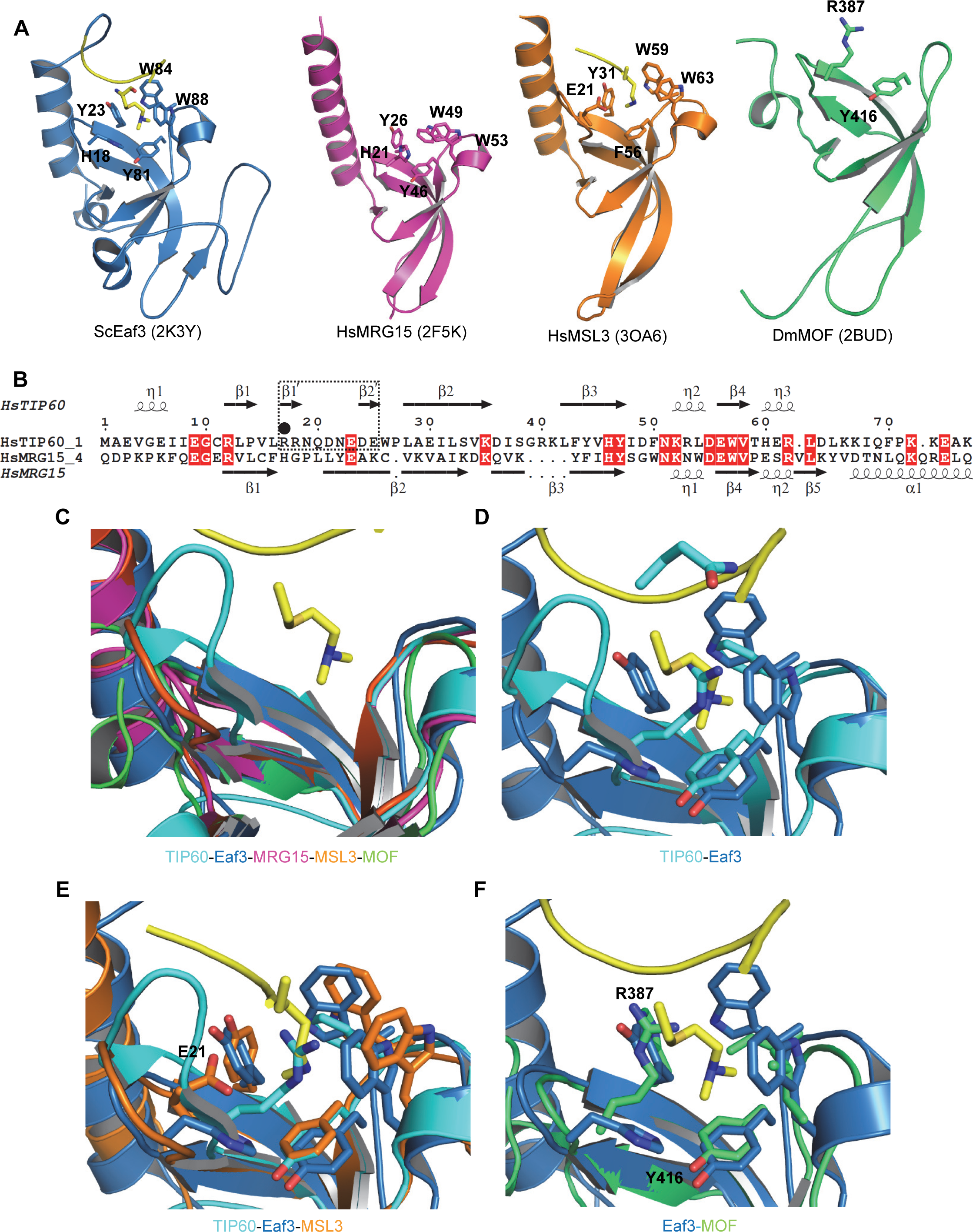
Structural comparison of chromo barrel domains. **(A)** Structure of selected chromo barrel domains: ScEaf3 (PDB code: 2K3Y, sky blue), MRG15 (PDB code: 2F5K, pink), MSL3 (PDB code: 3OA6, golden) and DmMOF (PDB code: 2BUD, light green). **(B)** Structure-based sequence alignment of chromo barrel domains of TIP60 and MRG15. Secondary structure elements of TIP60 and MRG15 are indicated above and below the sequence alignment, respectively. Black dash line box, the unique β harpin of TIP60-CB; black dot, the positive charged residues lining the hydrophobic cage of chromo barrel domains. The alignments were constructed with ClustalW[56] and refined with ESPript[57]. **(C-F)** Superimposition of the selected chromo barrel domain (C), TIP60 with ScEaf3 (D), MSL with TIP60 and ScEaf3 (E), DmMOF with ScEaf3 (F). Abbreviations: Hs, *Homo sapiens*; Sc, *Saccharomyces cerevisiae*; Dm, *Drosophila melanogaster*.

Structural comparison with other chromo barrel domains reveals that all the known histone binding chromo barrel domains (ScEaf3, MRG15 and MRG2) have a short loop in the potential histone binding site except MSL3, which has a similar long loop to the β hairpin of TIP60 in its chromo barrel domain (Fig. 3C and 3E). However, this loop does not block the histone binding site and the corresponding residue to Arg17 in MSL3 is Glu21, which is located outside the aromatic cage. In contrast to the positive charged residue, which clashes with the same charged methylated residue, the negative charged residue Glu21 might favor the methylated residue. Compared to the methyl-lysine complex structure of the chromo barrel domain of ScEaf3, the chromo barrel domain of DmMOF holds an incomplete cage, formed by only one aromatic residue Tyr416, and this incomplete cage is also occupied by a positive charged residue, Arg387 (Fig. 3F). These two features of DmMOF prevent the binding of methyl-lysine histone peptides to DmMOF, as reported previously[25]. Taken together, our structural and binding analysis presented here, coupled with the previously published chromo barrel domain structures, reveals that not all of the chromo barrel domains have the histone binding ability.

Whereas the position of phosphorylation target Tyr44 inside the amino acid sequence implies proximity to the aromatic cage, both solution and crystal structures indicate that the side chain points away from the aromatic cage, consistent with the residue’s position on the same β strand as cage residue Tyr47 (Fig. 2F). Mechanistic insight into the interplay between phosphorylation and putative peptide binding[2] remains elusive.

### Structural Comparison of the chromo barrel domain of TIP60 to other Royal family members

The chromo barrel domain reported here is also often called chromodomain or chromodomain-like domain or Tudor-knot domain in the protein domain annotation databases, such as SMART or InterPro, which indicates that the chromo barrel domain also belongs to Royal family domains, made up by chromodomain, Tudor domain, PWWP (Pro-Trp-Trp-Pro) domain and MBT (malignant brain tumor) repeat[28]. Sequence alignment of some chromo barrel domains with the typical chromodomain protein HP1, Tudor domain protein SMN and MBT repeat protein L3MBTL1 shows that the chromo barrel domain has the highest similarity to the chromodomain, especially for the part of the chrombox homology motif that is used to distinguish the chromodomain form other Royal family members[29] (Fig. 4A). A large number of Royal family domain-containing proteins have been reported as methylated-lysine and methylated-arginine readers[20,30]. These proteins use a hydrophobic cage, which is formed by 2-4 aromatic residues, to recognize and bind to the methylated lysine or arginine[19,20,31,32]. Sequence alignment indicates that most cage forming aromatic residues are highly conserved in Royal family domains (Fig. 4A). Normally, the histone methylation binding aromatic cage consists of 2-4 aromatic and long hydrophobic side chain residues with the cage being accessible to the methyl-lysine or methyl-arginine residue. Based on whether a complete and fully functional aromatic cage is present or not, the chromo barrel domain is divided into MOF-like and MSL3-like subgroups, which may bind to RNA and methylated residues, respectively[25]. According to this classification, TIP60 belongs to the MOF-like subgroup, which may not bind to the methylated residues and is consistent with our binding data (Fig. 1).

**Figure 4.**
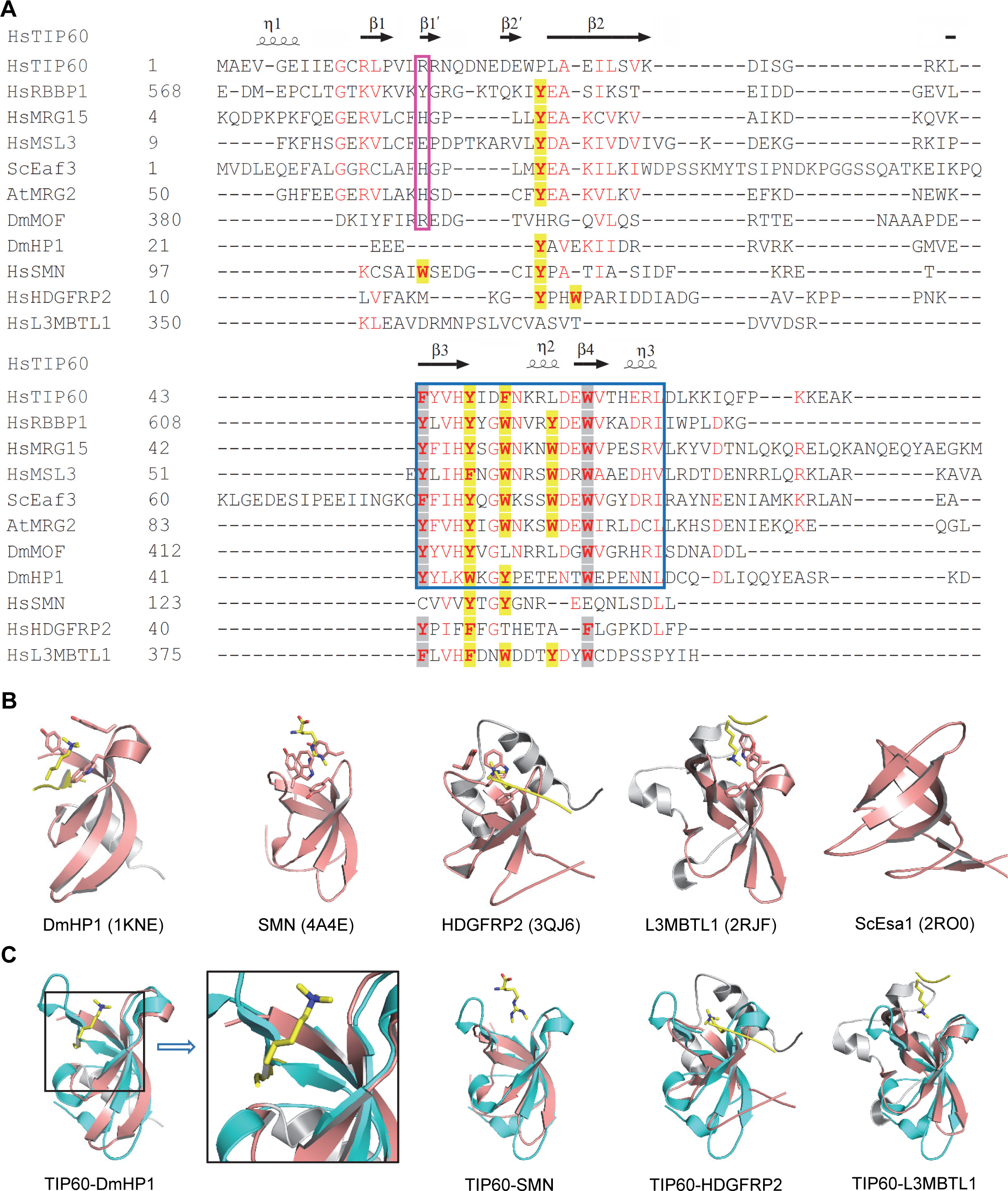
Sequence and structural comparison of Royal family domains. **(A)** Structure-based sequence alignment of selected chromo barrel domains and representative Royal family domains. Secondary structure elements of TIP60 are indicated above the sequence alignment. Yellow highlight, aromatic residues for cage forming; gray highlight, aromatic residues conserved in Royal family; pink box, positive charged residues lining the hydrophobic cage of chromo barrel domains; blue box, chromobox homology motif. The alignments were constructed with ClustalW[56] and refined with ESPript[57]. **(B)** Structure of selective Royal family domains: DmHP1 chromodomain (PDB code: 1KNE), SMN Tudor domain (PDB code: 4A4E), HDGFRP2 PWWP domain (PDB code: 3QJ6), L3MBTL1 MBT repeat (PDB code: 2RJF) and ScEsa1 knotted-Tudor domain (PDB code: 2RO0). **(C)** Superimposition of TIP60-CB (cyan) with chromodomain of DmHP1, Tudor domain of 53BP1, PWWP domain of HDGFRP2 and MBT repeat of L3MBTL1 (salmon). The secondary elements are shown as cartoon with the cage forming residues shown as sticks and the histone peptides are colored in yellow with the methylated lysines shown as sticks.

Although the sequence alignment suggests high sequence similarity between the chromo barrel domain and Royal family domains, structural comparison shows several structural differences. The common structural core of a Royal family domain is a β-barrel-like fold[28]. The complete tertiary structure of a specific Royal family domain is formed by adjoining this common β-barrel-like fold with adjacent β-strands or α-helices. In general, chromo barrel domain is composed of 5 β-strands and one α-helix[25,26], while chromodomain is made up by 3 β-strands and one α-helix[33–36], Tudor domain harbors 4-5 β-strands[37], PWWP domain contains 5 β-strands and 1 to 6 α-helices[38], and MBT, which forms a stable structure unit by using 2-4 repeats, consists of 5 β-strands and an extended α-helix arm from its neighbor (Fig. 3A and 4B)[18,39]. The Tudor-knot is named from the *Saccharomyces cerevisiae* Esa1 knotted Tudor domain, which is formed by core Tudor domain with extended N- and C-termini forming an antiparallel β-sheet and acting as a knot to help the Tudor domain to interact with RNA[40]. So the Tudor-knot is a special Tudor domain (Fig. 4B). The chromo barrel domain is also referred to as noncanonical chromodomain because it normally has two more beta strands than the canonical chromodomain, which just has 3 beta strands and 1 helix[29]. In addition, although both chromo barrel domain and chromodomain share one common structural feature, one α-helix, the α-helix of chromo barrel domain is packed against β-barrel core in different direction as chromodomain does (Fig. 3A and 4B). Although there are several structural difference, the β-barrel core of chromo barrel domain can superimpose very well with the other Royal family members, which also indicates the chromo barrel domain is a member of Royal family (Fig. 4C). Structural comparison also indicates that the extra beta strand (β1) takes over the histone peptide binding groove of typical chromodomain, which may explain the weak binding of histone peptide to chromo barrel domain (Fig. 4C).

In conclusion, our quantitative FP and ITC binding assays fail to identify any direct interaction between TIP60 and histone peptides. Whereas the crystal structure of TIP60-CB indicates that the formation of an aromatic cage is possible, it also suggests that access to the cage may be occluded by a basic amino acid side chain within a unique β hairpin. This observation alone does not rule out peptide binding, as such obstruction may be transient.

## Materials and methods

### Protein expression and purification

The chromo barrel domain of TIP60 (residues 1-80) was subcloned into a modified pET28-MHL vector. The encoded N-terminal His-tagged fusion protein was overexpressed in *Escherichia coli* BL21 (DE3) Codon plus RIL (Stratagene) cells at 15 °C and purified by affinity chromatography on Ni-nitrilotriacetate resin (Qiagen), followed by TEV protease treatment to remove the tag. Protein was further purified by Superdex75 gel-filtration (GE Healthcare, Piscataway, NJ). For crystallization experiments, purified protein was concentrated to 20 mg/mL in a buffer containing 20 mM Tris, pH 7.5, 150 mM NaCl and 1 mM DTT. The molecular weight of the purified protein was determined by mass spectrometry.

### Isothermal Titration Calorimetry (ITC)

All histone peptides were synthesized by Peptide 2.0 Inc. The concentrated proteins were diluted in 20 mM Tris, pH 7.5, and 150 mM NaCl. The lyophilized peptides were dissolved in the same buffer and pH was adjusted by adding NaOH. Peptide concentrations were estimated from the mass of lyophilized material. All measurements were performed at 25 ^0^C, using a VP-ITC microcalorimeter (GE Healthcare). Protein with a concentration of 50-100 μM was placed in the chamber, and the peptide with a concentration of 1-2 mM was injected in 25 successive injections with a spacing of 180 s and a reference power of 13 μcal/s. Control experiments were performed under identical conditions to determine the heat signals that arise from injection of the peptides into the buffer. Data were fitted using the single-site binding model within the Origin software package (MicroCal, Inc.).

### Fluorescence polarization assay

Peptides were synthesized and purified by Tufts University Core Services (Boston, MA, U.S.A.) with fluorescein-labeled C-termini. Binding assays were performed in 10 μL at a constant fluorescein labeled-peptide concentration of 40 nM and increasing amounts of protein at concentrations ranging from low to high micromolar in a buffer of 20 mM Tris, pH 7.5, 150 mM NaCl, 1 mM DTT, and 0.01% Triton X-100. All assays were performed in duplicate in 384-well plates, using the Synergy 2 microplate reader (BioTek) with an excitation wavelength of 485 nm and an emission wavelength of 528 nm. Data were corrected by background of the free labeled peptides and fitted to the ligand binding function using GraphPad Prism 5 software to determine the *Kd* values.

### Crystallization

Purified TIP60 (20 mg/mL) was mixed with trypsin at 1:1000 mass ratio[41] and crystallized using the sitting drop vapor diffusion method at 18 °C by mixing 0.5 μL of the protein with 0.5 μL of the reservoir solution. The crystals were obtained in a buffer containing 20% PEG 3350 and 0.2 M calcium acetate.

### Data collection and structure determination

Diffraction data were collected at low temperature at beam line 19ID of the Advanced Photon Source (Argonne, IL) and reduced with XDS[42] and AIMLESS[43] software. The structure was solved by molecular replacement with the program MOLREP[44] and coordinates extracted and transformed by PHASER[45] from the TIP60 chromo barrel domain solution structure (PDB code: 2EKO). The model was automatically retraced with BUCCANEER[46] program using PARROT[47] modified phases. The present model was obtained through iterative manual re-building with COOT[48], restrained refinement under “local” non-crystallographic symmetry restraints with REFMAC[49] and model validation with MOLPROBITY[50]. CCP4[51], PHENIX[52], PDB_EXTRACT[53] programs and the IOTBX library[54] were used in preparing model summaries (Table 1) and Protein Data Bank deposition.

### Accession number

Coordinates and structure factor amplitudes of TIP60-CB were deposited in the PDB with accession code 4QQG.

## Acknowledgements

We thank Wolfram Tempel for data collection, structure determination, and critical reading and editing of this manuscript, Amy K. Wernimont for reviewing a preliminary crystallographic model of the TIP60-CB. Results shown in this report are derived from work performed at Argonne National Laboratory, Structural Biology Center at the Advanced Photon Source. Argonne is operated by UChicago Argonne, LLC, for the U.S. Department of Energy, Office of Biological and Environmental Research under contract DE-AC02-06CH11357. This study was supported by the National Natural Science Foundation of China [grant number 31500613] and Hubei Chenguang Talented Youth Development Foundation. The SGC is a registered charity (number 1097737) that receives funds from AbbVie, Bayer Pharma AG, Boehringer Ingelheim, Canada Foundation for Innovation, Eshelman Institute for Innovation, Genome Canada through Ontario Genomics Institute [OGI-055], Innovative Medicines Initiative (EU/EFPIA) [ULTRA-DD grant no. 115766], Janssen, Merck KGaA, Darmstadt, Germany, MSD, Novartis Pharma AG, Ontario Ministry of Research, Innovation and Science (MRIS), Pfizer, Säo Paulo Research Foundation-FAPESP, Takeda, and Wellcome.

## Author contributions

Yanli Liu and Yuzhe Zhang purified and crystallized the protein; Ming Lei conducted the ITC assays; Xiao Liang conducted the FP assays; Peter Loppnau and Yanju Li cloned the constructs; Jinrong Min and Yanli Liu conceived and designed the study and wrote the paper. All authors analyzed the data and approved the final version of the manuscript.

